# AGO4 is specifically required for heterochromatic siRNA accumulation at Pol V-dependent loci in *Arabidopsis thaliana*

**DOI:** 10.1101/078394

**Authors:** Feng Wang, Michael J. Axtell

## Abstract

Genome-wide characterization of *AGO4*-dependent siRNAs revealed that *AGO4* is required for the accumulation of a small subset of heterochromatic siRNAs in *Arabidopsis thaliana*. These *AGO4-*depdenent siRNAs are likely secondary het-siRNAs produced by a self-reinforcing loop of RdDM. Slicing-defective *AGO4* is unable to fully complement het-siRNA accumulation from an *ago4* mutant, demonstrating the critical role of AGO4 catalytic ability in het-siRNA accumulation.

Summary: 152; Introduction: 618; Results: 1291; Discussion: 1013; Experimental procedures: 881; Acknowledgements: 24; Figure legends: 568; Author contribution: 25; Conflict of interest: 13; Funding: 34; References: 1289

**Summary:** In plants, 24 nucleotide long heterochromatic siRNAs (het-siRNAs) transcriptionally regulate gene expression by RNA-directed DNA methylation (RdDM). The biogenesis of most het-siRNAs depends on the plant-specific RNA polymerase IV (Pol IV), and ARGONAUTE4 (AGO4) is a major het-siRNA effector protein. Through genome-wide analysis of sRNA-seq data sets, we found that *AGO4* is required for the accumulation of a small subset of het-siRNAs. The accumulation of *AGO4*-dependent het-siRNAs also requires several factors known to participate in the effector portion of the RdDM pathway, including RNA POLYMERASE V (POL V), DOMAINS REARRANGED METHYLTRANSFERASE 2 (DRM2) and SAWADEE HOMEODOMAIN HOMOLOG 1 (SHH1). Like many AGO proteins, AGO4 is an endonuclease that can ‘slice’ RNAs. We found that a slicing-defective AGO4 was unable to fully recover *AGO4-*dependent het-siRNA accumulation from *ago4* mutant plants. Collectively, our data suggest that *AGO4*-dependent siRNAs are secondary siRNAs dependent on the prior activity of the RdDM pathway at certain loci.

## Introduction

24 nucleotide (nt) heterochromatic small interfering RNAs (het-siRNAs) are usually loaded into ARGONAUTE4 (AGO4) to direct repressive chromatic modifications and subsequent transcriptional gene silencing via RNA-directed DNA Methylation (RdDM) (Zilberman *et al.*, 2003; Qi *et al.*, 2006). Het-siRNA-induced transcriptional silencing plays important roles in transposable element silencing, stress responses and genome stability (Law and Jacobsen, 2010; Matzke and Mosher, 2014). The production of het-siRNAs in *Arabidopsis thaliana* usually requires the plant-specific RNA POLYMERASE IV (Pol IV) (Onodera *et al.*, 2005; Herr *et al.*, 2005; Blevins *et al.*, 2015; Zhai *et al.*, 2015), RNA-DEPENDENT RNA POLYMERASE 2 (RDR2) (Xie *et al.*, 2004; Kasschau *et al.*, 2007) and one or more DICER-LIKE (DCL) proteins (most predominantly DCL3; (Henderson *et al.*, 2006). A second plant-specific RNA polymerase, Pol V, generates scaffold RNAs targeted by het-siRNAs associated with AGO4 (Wierzbicki *et al.*, 2008; Wierzbicki *et al.*, 2009). This targeting is thought to recruit the *de novo* DNA methyltranserase DOMAINS REARRANGED 2 (DRM2) to the local chromatin, which acts to catalyze 5-methylation of cytosines (Cao and Jacobsen, 2002; Zhong *et al.*, 2014). The SAWADEE HOMEODOMAIN HOMOLOG 1 (SHH1) protein interacts with chromatin at Pol V transcribed loci, and recruits Pol IV to promote further siRNA biogenesis specifically from Pol V-dependent regions (Law *et al.*, 2013; H., Zhang *et al.*, 2013). Despite their positions at the effector portion of the RdDM pathway, Pol V and DRM2 are required for the accumulation of a subset of Pol IV-dependent het-siRNAs in *Arabidopsis* (Pontier *et al.*, 2005; Mosher *et al.*, 2008).

The *Arabidopsis* genome has 10 *AGO* genes. AGO4, AGO6, AGO8, and AGO9 form a monophyletic clade (Vaucheret, 2008; Mallory and Vaucheret, 2010; Fang and Qi, 2016). *AGO8* has been suggested as a pseudogene (Vaucheret, 2008). AGO4 and AGO6 both bind 24 nt het-siRNAs and contribute to the canonical RdDM pathway in a non-redundant fashion (Zheng *et al.*, 2007; Havecker *et al.*, 2010; Duan *et al.*, 2015). AGO6 also binds 21 nt siRNAs and act as a key effector of the non-canonical RDR6-RdDM pathway (McCue *et al.*, 2015; Panda *et al.*, 2016). AGO9, which is primarily expressed in female gametes, interacts with het-siRNAs and silence TEs in female gametes (Olmedo-Monfil *et al.*, 2010). Though AGO4, AGO6 and AGO9 are functionally related, the small RNA profile of an *ago4*/*ago6*/*ago9* triple mutant has not been reported yet.

According to the current model of the RNA-directed DNA methylation (RdDM) pathway, the biogenesis of het-siRNAs depends on Pol IV, RDR2, and primarily DCL3 (Law and Jacobsen, 2010; Matzke and Mosher, 2014), while AGO4 is not directly required for the biogenesis of het-siRNAs. However, the accumulation of certain het-siRNAs was shown to be dependent on AGO4 in previous reports (Qi *et al.*, 2006; Havecker *et al.*, 2010). It has been hypothesized that the accumulation of a subset of het-siRNAs depends on AGO4-mediated target slicing (Qi *et al.*, 2006). All 10 Arabidopsis AGOs have a conserved Asp-Asp-His (DDH) or Asp-Asp-Asp (DDD) motif thought to form a catalytic center for cleavage of target RNA. The target-slicing ability of AGO1 and AGO7 has been confirmed *in vivo* (Vaucheret, 2008; Fang and Qi, 2016). AGO10 can slice miRNA target *in vitro*, but it is still unclear if the slicer-activity is required for its function in plants (Ji *et al.*, 2011; Zhu *et al.*, 2011). AGO4, which specifically binds 24nt het-siRNAs, can slice synthetic het-siRNA targets *in vitro* (Qi *et al.*, 2006) as well as the passenger-strand of het-siRNA duplexes *in vivo* (Ye *et al.*, 2012). The *in vitro* and/or *in vivo* slicing ability of AGO4 is abolished by mutagenesis of the presumed catalytic triad (Qi *et al.*, 2006; Ye *et al.*, 2012). However, the genome-wide effects of AGO4 slicing on global small RNA accumulation have not been previously reported.

## Results

### Accumulation of a subset of 24 nt het-siRNAs depends on *AGO4* in *Arabidopsis*

To systematically study the profile of *AGO4*-dependent het-siRNAs and the effect of AGO4 catalytic activity on het-siRNA accumulation, we expressed wild-type *AGO4* (*pAGO4:FLAG-AGO4-DDH*, wt*AGO4* hereafter) or slicing-defective *AGO4* (*pAGO4:FLAG-AGO4-DAH*, *D742A* hereafter) driven by the native *AGO4* promoter in both the *ago4-4* single mutant background and the *ago4-4/6-2/9-1* triple mutant background in *Arabidopsis* (Fig S1a). Three T3 transgenic plants with comparable levels of protein accumulation (Fig S1b) were used to prepare three biological replicate sRNA-seq libraries. It is worth noting that *ago4-4* is in the Ws background, while *ago6-2* and *ago9-1* are in the Col-0 background. We therefore prepared three replicate control sRNA-seq libraries from both Ws and Col-0. We merged sRNA-seq libraries from the same genotype and aligned them to reference genome to study the overall small RNA size distribution in tested samples. Loci dominated by 24 nt small RNAs were the most abundant in all tested genotypes, and the fractions of small RNA from 24 nt small RNA-dominated loci were similar across different genotypes (Fig S2). sRNA-seq libraries from all backgrounds were then merged, aligned to the reference genome, followed by *de novo* definition of expressed small RNA clusters. The 24 nt siRNA clusters that were *de novo* annotated are listed in Data S1.

We first examined siRNA accumulation in the Ws background, to compare *ago4-4* to the wild-type. A differential expression analysis was performed by comparing raw read counts from our *de novo* annotated small RNA loci for all libraries in Ws background. A principal component analysis (PCA) plot was prepared to visualize the overall differences between samples (Fig 1a). The biological replicates were grouped together, indicating good reproducibility (Fig 1a). *ago4-4*/*wtAGO4* grouped closely with the Ws wild-type, suggesting complementation of small RNA accumulation by expression of *wtAGO4* in the *ago4-4* background (Fig 1a). *ago4-4* and *ago4-4*/*D742A* were distinct from each other and from the wild-type and *ago4-4/wtAGO4* genotypes (Fig 1a).

**Figure 1.**
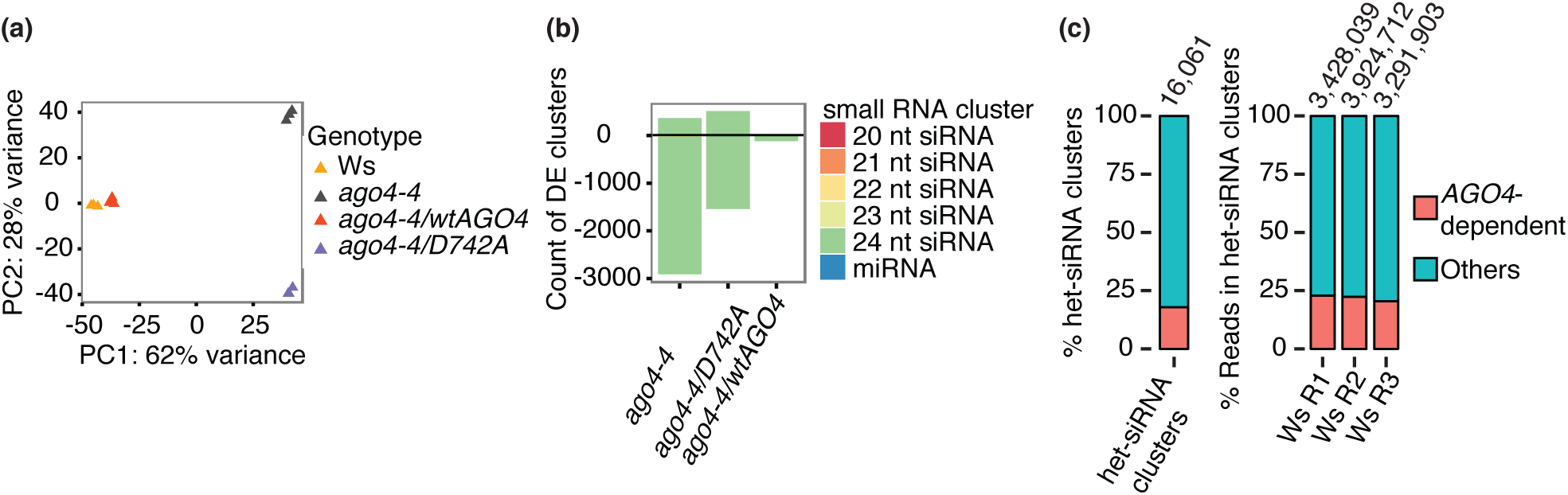
Identification of *AGO4*-dependent small RNA clusters in *Arabidopsis thaliana*. (a) Principal component analysis demonstrating overall relationships between sRNA-seq libraries in Ws background. (b) Number of differentially expressed (DE) clusters in the indicated genotypes and small RNA clusters compared with Ws wild type. DE clusters were defined as clusters with at least 2-fold change compared to wild type at a false discovery rate of 1%. (c) Percentage of *AGO4*-dependent clusters and *AGO4*-dependent small RNAs in 24 nt siRNA loci. *AGO4*-dependent clusters were defined as clusters with at least 2-fold less accumulation in *ago4-4* compared with Ws.

Most differentially accumulated clusters were dominated by 24 nt siRNAs (Fig 1b). In *ago4-4*, 2,912 clusters were down-regulated relative to wild-type; we defined these as *AGO4*-dependent siRNA clusters (Fig 1b). Most of these (2,879) were dominated by 24 nt siRNAs. In contrast, only 121 clusters were down regulated in *ago4-4*/*wtAGO4*, indicating nearly full complementation of small RNA accumulation by *wtAGO4* (Fig 1b). Intriguingly, an intermediate amount of clusters (1,541, Fig 1b) was down-regulated in *ago4-4*/*D742A*, which suggested that slicing-defective AGO4 partially recovers the accumulation of *AGO4*-dependent small RNAs. Most 24 nt-dominated siRNA clusters are not *AGO4*-dependent. Only about 18% of the *de novo* annotated 24 nt siRNA clusters, which contained about 22% of small RNAs in Ws wild-type, were dependent on *AGO4* (Fig 1c).

### Accumulation of small RNAs in *AGO4*-dependent clusters requires *NRPE1*, *DRM2* and *SHH1*

We classified the 16,061 *de novo* annotated 24 nt siRNA-dominated clusters into different groups based on *AGO4*-dependency (Data S1). As stated above, 2,879 24 nt-dominated siRNA clusters were *AGO4*-dependent (FDR=0.01). We found another 1,359 24 nt-dominated clusters that were clearly *AGO4-*independent (FDR=0.01). Another 354 24 nt-dominated clusters were up-regulated in *ago4-4* (FDR=0.01), and the *AGO4-*dependency of the remaining 11,469 24 nt-dominated siRNA clusters could not be reliably inferred using our strict statistical tests, primarily due to low expression levels. We analyzed sRNA-seq accumulation from the *AGO4*-dependent and *AGO4*-independent clusters using data from *nrpd1-4*, *nrpe1-12*, *drm2-2*, and *shh1-1* mutants (Law *et al.*, 2013), using accumulation of clusters overlapping high-confidence *MIRNA* loci (Kozomara and Griffiths-Jones, 2014) as a control (Fig 2a). Note that *NRPD1* and *NRPE1* encode the catalytic sub-units of Pol IV and Pol V, respectively. In *nrpd1*, siRNA accumulation was strongly down-regulated in both *AGO4*-dependent and *AGO4*-independent clusters (Fig 2a). In contrast, *AGO4*-dependent clusters were much more strongly affected in the *nrpe1*, *drm2*, and *shh1* backgrounds compared to AGO4-independent clusters (Fig 2a). We then normalized small RNA accumulation in *AGO4*-dependent and *AGO4*-independent het-siRNA clusters based on wild-type plants. We observed significantly reduced small RNA accumulation (Mann-Whitney test, p<0.01) in *AGO4*-dependent clusters relative to *AGO4*-independent clusters in all analyzed RdDM mutants except *nrpd1* (Fig 2b). Using small RNA-seq data from *nrpe1-1* plants (Lee *et al.*, 2012), we defined 2,827 *NRPE1*-dependent small RNA clusters, the majority of which overlapped *AGO4*-dependent siRNA clusters (Fig 2c). This extent of overlap far exceeded the number expected by random chance (Fig 2d). Collectively, these data indicate that the subset of 24 nt dominated siRNA loci that depend on *AGO4* for accumulation are those that are also dependent on *NRPE1*, *DRM2*, and *SHH1*.

**Figure 2.**
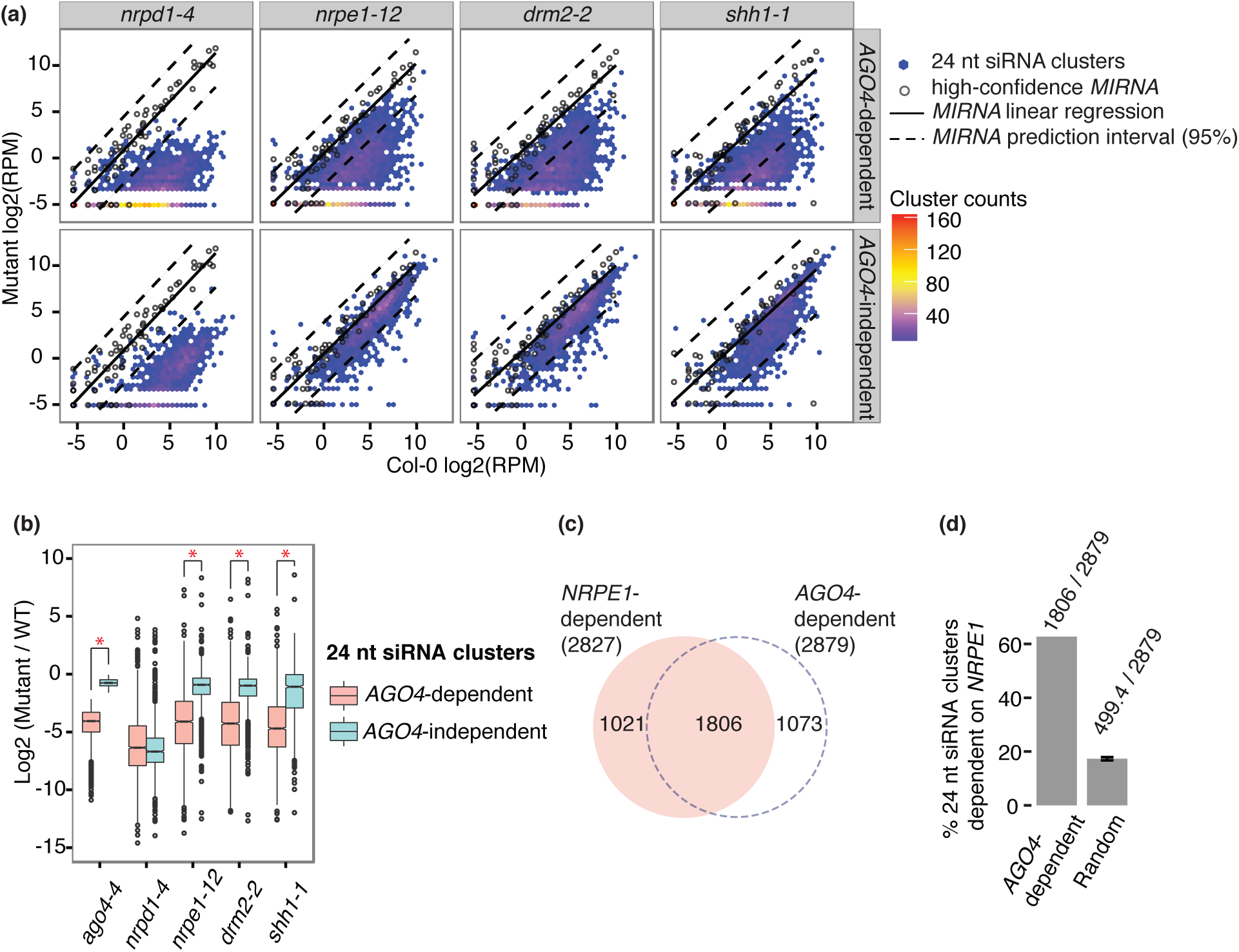
*AGO4*-dependent and *AGO4*-independent 24 nt siRNA clusters in other RdDM mutants. (a) Small RNA accumulation from *AGO4*-dependent and *AGO4*-independent 24 nt siRNA clusters in indicated RdDM mutants. A hexbin plot of Log2 transformed reads per million (RPM) in indicated geno-types were plotted. A linear regression (solid line) and associated 95% prediction interval (dashed lines) was plotted based upon accumulation from clusters overlapping high-confidence *MIRNA* loci. (b) Normalized small RNA accumulation in *AGO4*-dependent and *AGO4*-independent clusters in indicated RdDM mutants. Boxplots show medians (horizontal lines), the 1st-3rd quartile range (boxes), 95% confidence of medians (notches), other data out to 1.5 times the interquartile range (whiskers) and outliers (dots). Asterisks indicate significant differences (Mann-Whitney U test, p<0.01) between *AGO4*-dependent and *AGO4*-independent clusters in the indicated mutant. (c) Venn diagram showing the overlap of *AGO4*-dependent and *NRPE1*-dependent 24 nt siRNA clusters. (d) Percentage of overlap between *AGO4*-dependent and *NRPE1*-dependent clusters. Overlaps expected by random chance were estimated by by randomly choosing 2827 and 2879 clusters from all 24 nt siRNA clusters. The mean and standard deviation (n=10) of randomly overlapping percentages are shown.

### An AGO4 catalytic residue is required for full accumulation of most *AGO4-*dependent 24 nt siRNAs

We then compared complementation of siRNA accumulation from *AGO4*-dependent clusters between the *wtAGO4* and *AGO4-D742A* transgenic lines. Small RNA accumulation was recovered in *wtAGO4* from nearly all *AGO4*-dependent clusters, but only from a small subset of loci in the slicing-defective *AGO4-D742A* plants (Fig 3a). We defined *AGO4-D742A* complemented loci as those that were significantly down-regulated in the *ago4-4* background but not in the *ago4-4/AGO4-D742A* transgenic plants (Fig 3b). Conversely, *AGO4-D742A* non-complemented loci were defined as those that were significantly down-regulated in both *ago4-4* and *ago4-4/AGO4-D742A* (Fig 3b). By this measure, half (49.9%) of the *AGO4-*dependent siRNA loci required AGO4 catalytic activity for their accumulation. Detailed examination of accumulation levels revealed that recovery was generally not to full wild-type levels at loci designated as complemented by *AGO4-D742A* (Fig 3c). We conclude that that the catalytic ability of AGO4 is important for full accumulation of most *AGO4*-dependent 24 nt siRNAs, but to varying degrees at different loci.

**Figure 3.**
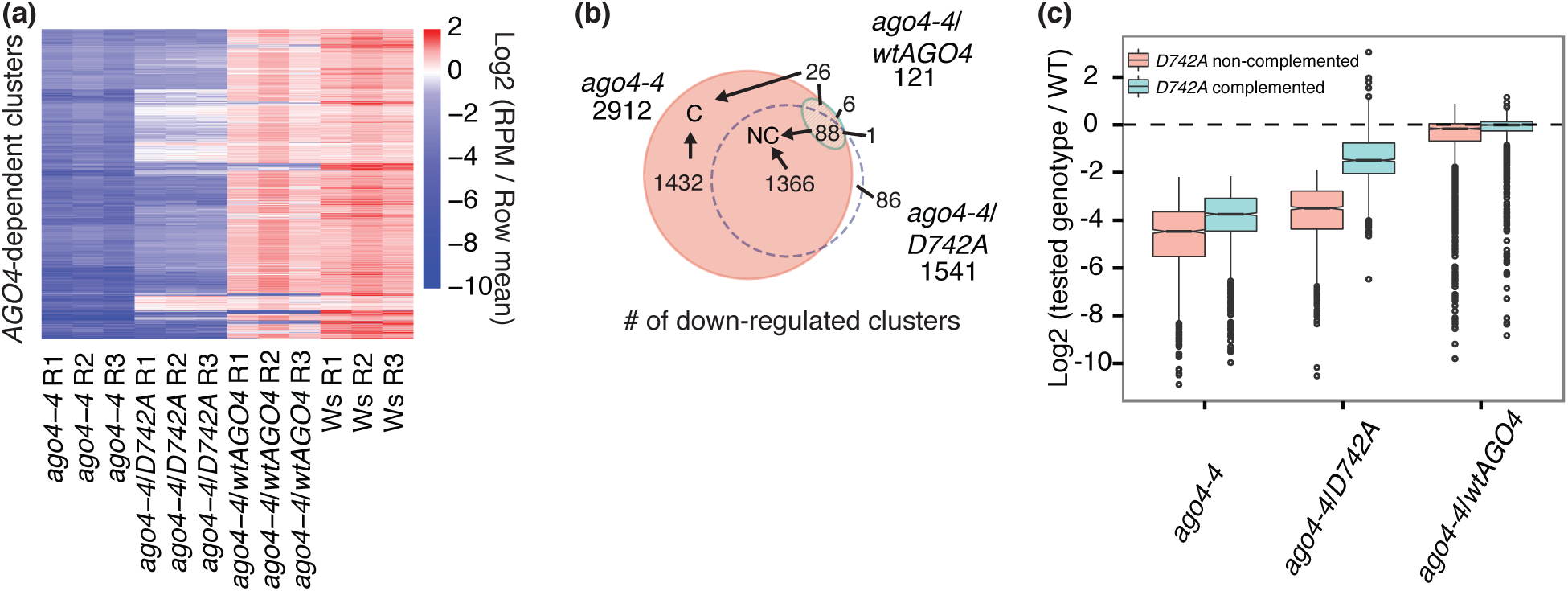
Slicing-defective *AGO4-D742A* partially complements small RNA accumulation from *AGO4*-dependent siRNA loci. (a) Heatmap showing normalized (log2-transformed and mean-centered) small RNA accumulation from *AGO4*-dependent clusters in indicated genotypes and replicates. (b) Euler diagram showing overlaps between down-regulated small RNA clusters (FDR=0.01) in the indicated genotypes compared to Ws wild-type. C: Complemented by the *AGO4-D742A* transgene; NC: Not complemented by the *AGO4-D742A* transgene. (c) Normalized small RNA accumulation levels from *AGO4*-dependent loci that were complemented or not complemented by the *AGO4-D742A* transgene. The ratio of small RNA accumulation in indicated geno-types over that in the Ws wild-type was computed and then log2-transformed. Boxplots show medians (horizontal lines), the 1st-3rd quartile range (boxes), 95% confidence of medians (notches), other data out to 1.5 times the interquartile range (whiskers) and outliers (dots).

### Slicing-defective AGO4 partially complements small RNA accumulation in the *ago4-4/ago6-2/ago9-1* triple mutant

*AGO4*, *AGO6*, and *AGO9* have related but non-redundant functions in gene silencing, and all three can bind 24 nt siRNAs (Havecker *et al.*, 2010). We obtained the triple mutant *ago4-4/ago6-2/ago9-1* and analyzed small RNA expression levels from inflorescence tissue. Significant ecotype-specific changes in small RNA accumulation levels were observed between Ws (the parental background of the *ago4-4* allele) and Col-0 (the parental background of the *ago6-2* and *ago9-1* alleles) (Fig S3a). About 15% of the small RNA clusters had significant differential accumulation (FDR = 0.01) when comparing Ws and Col-0 (Fig S3b). Different small RNA accumulation in these DE clusters was presumably caused by the different genetic backgrounds. We therefore excluded these loci from our analyses.

When analyzing the remaining, ecotype-insensitive small RNA clusters, we observed that the Col-0, Ws, and *ago4-4*/*wtAGO4* samples were tightly grouped (Fig 4a). This demonstrates both the effective removal of clusters that have ecotype-specific differences in accumulation, as well as strong complementation by the *wtAGO4* transgene. While *ago4-4*/*6-2*/*9-1*/*wtAGO4* strongly diverged from *ago4-4*/*6-2*/*9-1*, *ago4-4*/*6-2*/*9-1*/*D742A* showed only minimal differences from *ago4-4*/*6-2*/*9-1* (Fig 4a). This implies that introduction of wild-type *AGO4,* but not a slicing-defective *AGO4,* can rescue much of the small RNA accumulation defects of the triple mutant. Full elimination of AGO4-clade AGOs didn’t affect accumulation of the majority of 24 nt siRNA clusters: About 22% (3005/13602) of the 24 nt siRNA clusters in Col-0 were *AGO4*/*AGO6*/*AGO9*-dependent, and these clusters contributed only about 15% of the small RNA reads (Fig 4b). Only about 24% (719/3005) of the *AGO4/AGO6/AGO9-*dependent clusters were not complemented by *wtAGO4* (Fig 4c), indicating that *AGO6* and/or *AGO9* are required for accumulation from relatively few clusters. Similar to the single-mutant analysis (Fig 3), many of the *AGO4/AGO6/AGO9-*dependent clusters were not complemented by *AGO4-D742A* (Fig 4c). In addition, even the set of loci that were designated as complemented by *AGO4-D742A* still generally showed less accumulation then observed with the *wtAGO4* transgene (Fig 4d). Overall, these analyses demonstrate that *AGO4* is required for the accumulation of a much larger number of siRNAs compared to *AGO6* and *AGO9* in inflorescences, and that the slicing activity of AGO4 is required for full accumulation of most of these siRNAs.

**Figure 4.**
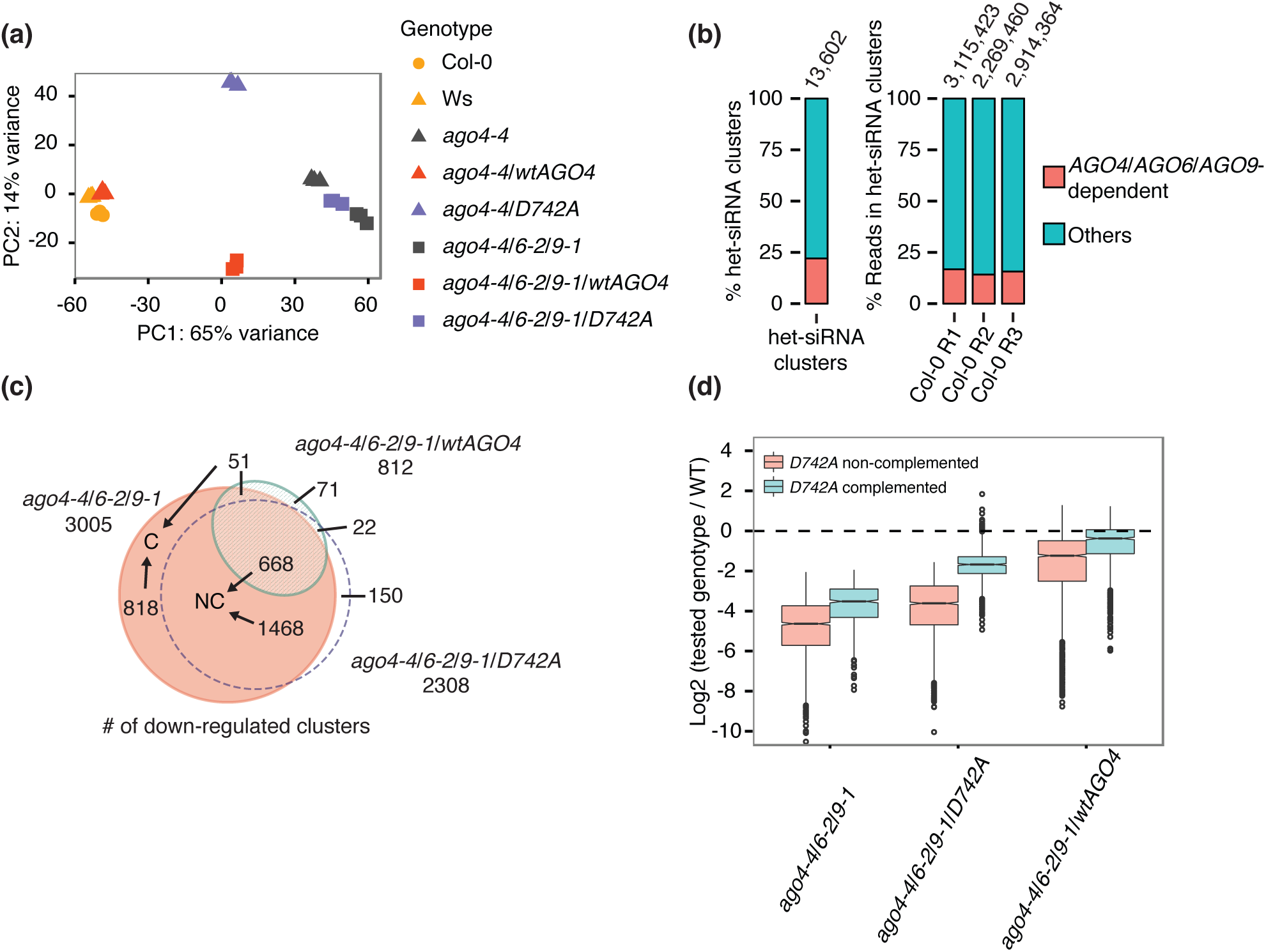
Slicing-defective *AGO4-D742A* partially complements small RNA accumulation in the *ago4-4*/*6-2*/*9-1* background. (a) Principal component analysis demonstrating overall relationships between all sRNA-seq libraries from ecotype-insensitive small RNA clusters. The first two principal components were shown for tested samples. (b) Percentage of *AGO4*/*AGO6*/*AGO9*-dependent clusters and *AGO4*/*AGO6*/*AGO9*-dependent small RNAs in 24 nt siRNA loci. *AGO4*/*AGO6*/*AGO9*-dependent clusters were defined as clusters with at least 2-fold less accumulation in *ago4-4*/*ago6-2*/*ago9-1* compared with Col-0. (c) Euler diagram showing the number of significantly down-regulated small RNA clusters (FDR=0.01) in indicated genotypes compared with Ws wild-type. C: *AGO4-D742A* complemented clusters in *ago4-4*/*6-2*/*9-1* background; NC: *AGO4-D742A* non-complemented clusters in *ago4-4*/*6-2*/*9-1* background. (d) Normalized small RNA accumulation in *AGO4-D742A* complemented and non-complemented clusters. The ratio of small RNA accumulation in indicated genotypes over that in the Ws wild-type was computed and then log2-transformed. Boxplots show medians (horizontal lines), the 1st-3rd quartile range (boxes), 95% confidence of medians (notches), other data out to 1.5 times the interquartile range (whiskers) and outliers (dots).

## Discussion

### Most 24 nt siRNAs do not require *AGO4, AGO6*, *or AGO9* for accumulation

*AGO4* is required for 24 nt small RNA at some loci, but not others (Zilberman *et al.*, 2003; Qi *et al.*, 2006). Our genome-wide analysis confirms this observation, and quantifies the extent of the dichotomy: Most 24 nt siRNA loci are unaffected by loss of *AGO4*, while only a small subset have siRNA accumulation defects. Even when all three functional members of the *AGO4* clade (Vaucheret, 2008) are removed in the *ago4-4/6-2/9-1* triple mutant, accumulation of 24 nt siRNAs from most loci is unaffected. This situation seems to contrast to the relationship between the major *Arabidopsis* miRNA-binding Argonaute AGO1 and miRNAs: In the null mutant *ago1-3*, accumulation of the majority of miRNAs is decreased (Vaucheret *et al.*, 2004; Arribas-Hernández, Kielpinski, *et al.*, 2016). Why might the majority of 24 nt siRNAs maintain stable accumulation levels in the absence of AGO4, AGO6, and AGO9? One possibility is that they are stabilized by AGO3. Despite not being a member of the AGO4 clade, *Arabidopsis* AGO3 is primarily associated with 24 nt siRNAs, and can partially complement the DNA methylation defects seen in the *ago4* mutant (Z., Zhang *et al.*, 2016). Alternatively, many 24 nt siRNAs might be stabilized by association with non-AGO RNA binding proteins, or perhaps not require protein binding at all.

### *AGO4-*dependent siRNAs are likely secondary siRNAs

Two models have been proposed to explain why some 24 nt siRNAs are dependent on *AGO4*. Qi *et al*. (2006) hypothesized that *AGO4-*dependent siRNAs might reflect target slicing-dependent secondary siRNA biogenesis similar to that which is sometimes observed from miRNA targets (Fei *et al.*, 2013). In this model, double-stranded RNA could be synthesized from using AGO4-sliced primary transcripts, which are then further processed into 24 nt secondary siRNAs by DCL3. Because Pol V makes chromatin-associated, long non coding RNAs that are targeted by AGO4 (Wierzbicki, 2012), the sliced-secondary siRNA model predicts that *AGO4-*dependent siRNAs would also be *NRPE1*-dependent. Our analysis shows that this prediction is supported by the data: most *NRPE1*-dependent siRNA clusters are also *AGO4-*dependent, and *vice-versa*. However, we also found that *AGO4-*dependent siRNAs also tend to be *DRM2-*dependent. This isn’t an obvious prediction of the sliced-secondary siRNA model because DRM2, a *de novo* DNA methyltransferase, is thought to be recruited to chromatin in the vicinity of an AGO4-Pol V interaction. An alternative model proposed that an initial wave of *de novo* AGO4/Pol V-dependent DNA methylation at a locus could subsequently recruit Pol IV and thus produce secondary siRNAs in a self-reinforcing loop (Pontier *et al.*, 2005). Our observation that *ago4*, *nrpe1*, *drm2,* and *shh1* were all required for accumulation of the same subsets of 24 nt siRNA loci is fully consistent with the self-reinforcing loop model for secondary het-siRNAs. Intriguingly, much stronger reduction of het-siRNA accumulation was observed in *nrpd1-4* than in *shh1-1*, suggesting that SHH1 may be specifically required for guiding Pol IV to the regions targeted by AGO4-dependent, self-reinforcing silencing.

### On the role of AGO4-catalyzed slicing

A full description of the functions of AGO4-catalyzed endonuclease activity (*e.g.* slicing) remains elusive. In other systems, two general functions of AGO-catalyzed slicing have been described: Slicing of passenger strands during AGO-loading of a small RNA duplex (Matranga *et al.*, 2005), and slicing of target RNAs (Qi *et al.*, 2005). For *Arabidopsis* AGO1, both *in vitro* and *in vivo* experiments demonstrate that AGO1-catalyzed slicing is not required for miRNA loading, but is required for many aspects of target regulation (Iki *et al.*, 2010; Carbonell *et al.*, 2012; Arribas-Hernández, Kielpinski, *et al.*, 2016; Arribas-Hernández, Marchais, *et al.*, 2016). In contrast, *in vitro* and *in vivo* data have demonstrated that AGO4-catalyzed slicing is required for passenger strand removal during siRNA loading and subsequent nuclear localization of the AGO4-siRNA complex (Ye *et al.*, 2012). Although AGO4 can slice a free target RNA *in vitro* (Qi *et al.*, 2006), to our knowledge there is no direct evidence of AGO4-catalyzed slicing of Pol V target RNAs *in vivo*. Our analysis showed that the catalytic capability of AGO4 is critical for the full accumulation of nearly all *AGO4-*dependent siRNAs. Many siRNAs were not rescued at all by slicing-defective AGO4, and even those that showed some degree of complementation almost never recovered to the extent allowed by complementation with the wild-type AGO4. The dependency of *AGO4*-dependent siRNAs upon AGO4-catalyzed slicing could be fully explained by defects in siRNA loading (Ye *et al.*, 2012). In either the sliced-secondary siRNA or self-reinforcement secondary siRNA models, lack of proper loading and subsequent nuclear localization of the ‘primary’ siRNAs would prevent accumulation of the *AGO4*-dependent sub-population.

Ye *et al*. reported that passenger strand removal mediated by AGO4 slicing is required for nuclear location of AGO4 (Ye *et al.*, 2012). Why could any complementation occur at all in the slicing defective mutant *AGO4-D742A* in our study? One hypothesis is that the passenger strand removal for proper AGO4-loading may not be completely dependent on slicing. AGO1-mediated slicing is not required for the unwinding of miRNA/miRNA* duplexes during AGO1-loading (Iki *et al.*, 2010; Carbonell *et al.*, 2012; Arribas-Hernández, Kielpinski, *et al.*, 2016; Arribas-Hernández, Marchais, *et al.*, 2016). Slicing-independent miRNA loading may be efficient because of the mismatches and bulges in common in miRNA/miRNA* duplexes (Iki *et al.*, 2010). In the case of AGO4-loading, where siRNA duplexes are perfectly complementary, a slicing-independent mechanism might still contribute to passenger strand removal, but with a much lower efficiency.

Whether or not AGO4-catalyzed slicing occurs at the targeting stage (*e.g.* in the nucleus upon targeted Pol V transcripts) remains unclear. If so, it would seem to present difficulties for the current model of RdDM, which supposes that a stable tethering of AGO4-siRNA complexes to nascent RNAs is required to recruit DRM2 to the vicinity. Conversely, if slicing is not used at the targeting stage, the challenge becomes understanding how it is prevented *in vivo*, given that *in vitro* AGO4-siRNA complexes are perfectly competent to direct target cleavage (Qi *et al.*, 2006). Resolution of these questions is an important goal for the future that will further illuminate the mechanisms of RdDM.

### Experimental procedures

### Plant materials and growth condition

All *Arabidopsis thaliana* plants were grown at 21°C with 16 h light/8 h dark. *ago4-4* (FLAG_216G02) was from INRA T-DNA transformants in the Wassilevskija (Ws) ecotype. *ago6-2* (SALK_031553) and *ago9-1* (SALK_127358) were from Salk T-DNA transformants in the Columbia-0 (Col-0) ecotype. The *ago4-4/ago6-2/ago9-1*triple mutant was generated by crossing *ago4-4* to *ago6-2* first, and crossing the *ago4-4/6-2* double mutant to *ago9-1*. Homozygous mutants were selected by genotyping using primers that specifically amplify T-DNA inserted alleles. All the genotyping primers are listed in Table S1.

### Cloning of wild-type and slicing-defective AGO4

cDNA encoding *AGO4* (*AT2G27040*) was amplified from *Arabidopsis thaliana* cDNA in Col-0 ecotype. A *FLAG* tag was inserted at the 5’ of *AGO4* cDNA right after start codon by PCR. The *FLAG*-tagged *AGO4* sequence was sub-cloned into the pGII0179 vector. A ~ 2 kb DNA sequence located upstream of the start codon of *AGO4* in Col-0, and a ~ 500 bp DNA sequence downstream of stop codon of *AGO4* in Col-0, were further sub-cloned into *AGO4* expression vector as native promoter and terminator (*pAGO4:FLAG-AGO4*). Mutagenesis of the catalytic motif of *AGO4* was performed by overlapping extension PCR. Primers with desired changes, which encode alanine instead of aspartic acid at the 742th amino acid position of AGO4, were used to introduce slicing defective mutation. The wild-type *AGO4* sequence in *AGO4* expression vector was then swapped by mutagenized *AGO4* to generate slicing-defective *AGO4* expression vector (*pAGO4:FLAG-AGO4-D742A*). The hygromycin-B phosphotransferase gene was inserted into both wild-type and slicing-defective *AGO4* expression vectors for hygromycin resistance selection in transgenic plants. All primers used for subcloning are listed in Table S1.

### Plant transformation and transgenic plant selection

Wild-type or slicing-defective *AGO4* expression vector was introduced into *ago4-4* or the *ago4-4*/*ago6-2*/*ago9-1* background by floral dip with *Agrobacterium tumefaciens* strain *GV3101* bearing the pSOUP plasmid and designated expression vectors. Transgenic plants were selected on 1/2 strength Murashige-Skoog plates supplemented with 15mg/L Hygromycin-B. Independent transgenic lines with single insertion were selected in the T2 generation. Homozygous lines with comparable wild-type or slicing-defective AGO4 protein accumulation in the T3 generation were further selected to prepare sRNA-seq libraries.

### sRNA-seq library preparation

Libraries were constructed by using 1µg total RNA extracted from *Arabidopsis* immature inflorescence tissue as described in Wang *et al*. (2016). Three biological replicates from each genotype were prepared. Raw data have been deposited at NCBI GEO under accession number GSE79119 (Col-0 samples) and GSE87333 (all other samples). Details for sRNA-seq libraries are listed in Table S2.

### Differential expression analysis

sRNA-seq data sets, including libraries from wild-type AGO4 and slicing-defective AGO4 transgenic lines in *ago4-4* and *ago4-4/ago6-2/ago9-1* background, mutant controls of *ago4-4* and *ago4-4/ago6-2/9-1*, wild-type controls of Col-0 and Ws, were merged and run with ShortStack 3.3 (Johnson *et al.*, 2016) with options --adapter TGGAATTC --mincov 50. All sRNA-seq libraries were aligned to the *Arabidopsis* TAIR10 reference genome.

A matrix of raw read counts from *de novo* annotated small RNA clusters in all three biological replicates of different genotypes were used for differential expression analysis with the R package DESeq2 (Love *et al.*, 2014). Clusters with at least a 2-fold change relative wild-type at a 1% false discovery rate were defined as differentially expressed.

To identify differentially expressed clusters in *nrpe1* compared to Col-0, sRNA-seq data sets from a previous study (Lee *et al.*, 2012) with three biological replicates of *nrpe1-1* and three biological replicates of Col-0 were analyzed with the same pipeline as described above, except that small RNA clusters were previously annotated by analyzing the AGO4-related data sets. sRNA-seq libraries used in this analysis are listed in Table S2.

### Heatmap of small RNA accumulation in AGO4-dependent clusters

To generate the heatmap for small RNA accumulation visualization, we first transformed read per million (RPM) data in AGO4-dependent cluters with the equation E = log_2_(R_i_/R_m_), where E is the input for heatmap, R_i_ is the RPM of a cluster in a sRNA-seq library, R_m_ is the mean RPM of a cluster across different sRNA-seq libraries been analyzed for the heatmap. The matrix of transformed RPM was then used for heatmap preparation with the R package pheatmap (Kolde, 2015).

### Euler diagrams

All Euler diagrams in this study were prepared with eulerAPE 3.0 (Micallef and Rodgers, 2014).

### Small RNA accumulation in *nrpd1-4*, *nrpe1-12*, *drm2-2, shh1-1,* and *ago4-4*

sRNA-seq libraries from a study (Law *et al.*, 2013) containing samples from *nrpd1-4*, *nrpe1-12*, *drm2-2, shh1-1* and Col-0 were aligned to the *Arabidopsis* TAIR10 genome using ShortStack 3.3 (Johnson *et al.*, 2016) with a locifile specifying small RNA clusters which were defined in the AGO4 sRNA-seq data sets. The 3’ adapters were removed with the option --adapter TGGAATTC. Before log2 transformation, a value of 0.5 was added to all raw counts. Log2 transformed RPMs of 24 nt siRNA clusters from *nrpd1-4*, *nrpe1-12, drm2-2* and *shh1-1* as well as Col-0 were plotted to illustrate small RNA accumulation in 24 nt siRNA clusters. Log2 transformed RPMs of high-confidence miRNA genes were also plotted. The linear regression and 95% predicted intervals were calculated based on the distribution of high-confidence miRNA genes. Small RNA accumulation at 24 nt siRNA loci in indicated RdDM mutants was then normalized to corresponding wild-type plants, with equation N = log2 (RPM_mutant_/RPM_WT_). Statistical differences between AGO4-dependent and AGO4-independent clusters were tested using the Mann-Whitney U test.

## Author contributions

MJA conceived of the project. FW generated transgenic plants, constructed small RNA-seq libraries and performed data analysis. MJA and FW wrote the manuscript.

## Acknowledgements

The authors thank Penn State University Genomic Core Facility for small RNA-seq services, and all members of the Axtell Lab for constructive comments.

## Supporting information

**Figure S1.**
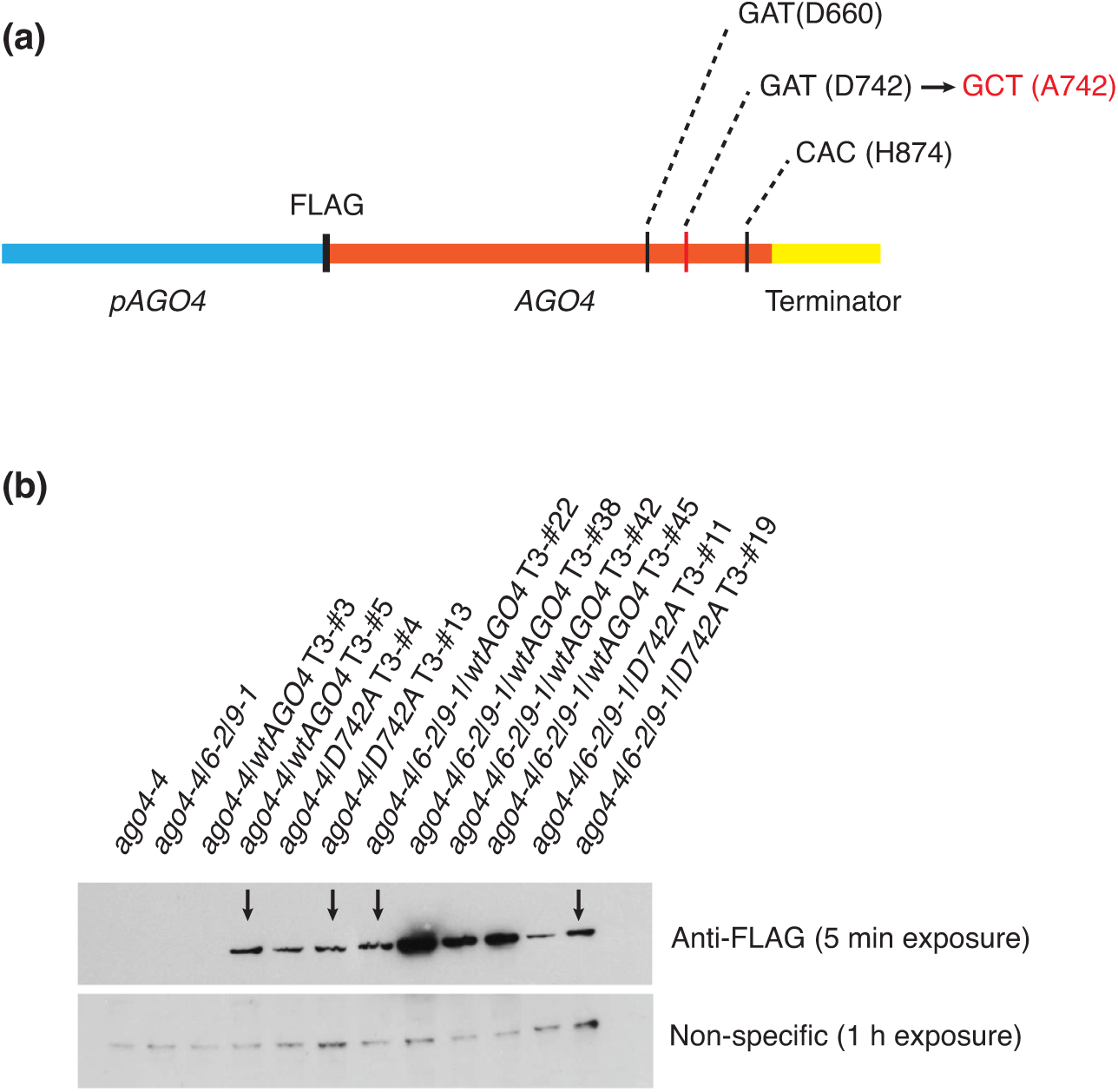
Expression of wild-type AGO4 and slicing-defective AGO4 proteins in transgenic plants. (a) Schematic of transgenes. Indicated codons correspond to the catalytic residues required for slicing. Codon color-coded by red represented the mutagenesis of codon 742. (b) Anti-FLAG immunoblot of FLAG-tagged AGO4 in T3 lines of the indicated transgenic plants. Transgenic lines that were chosen for sRNA-seq library preparation, based on approximately equal accumulation of AGO4 protein, are indicated by arrows.

**Figure S2.**
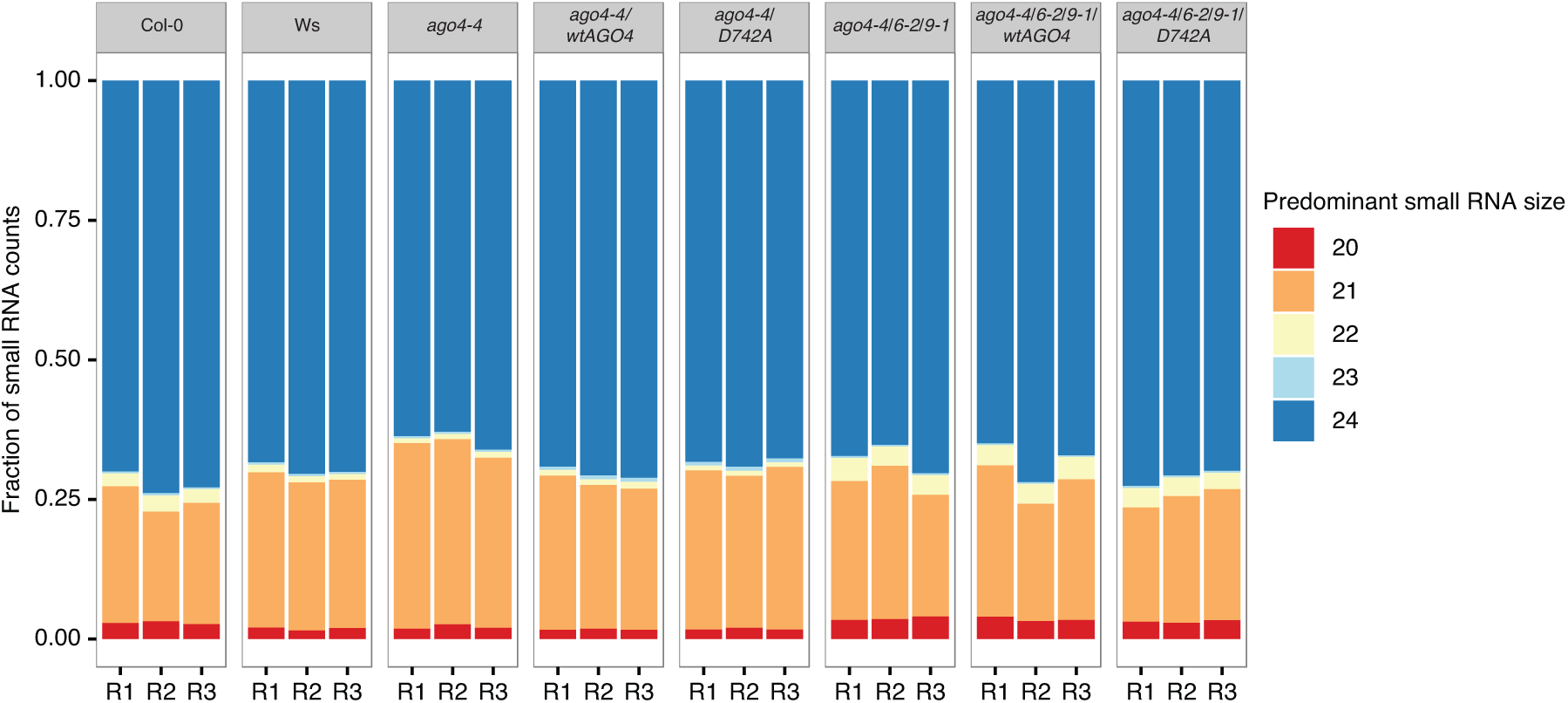
Overall size profiles of small RNAs in tested genotypes. Fractions of small RNA clusters with different predominant sizes in indicated genotypes are shown. R1, R2 and R3 represent three biological replicates.

**Figure S3.**
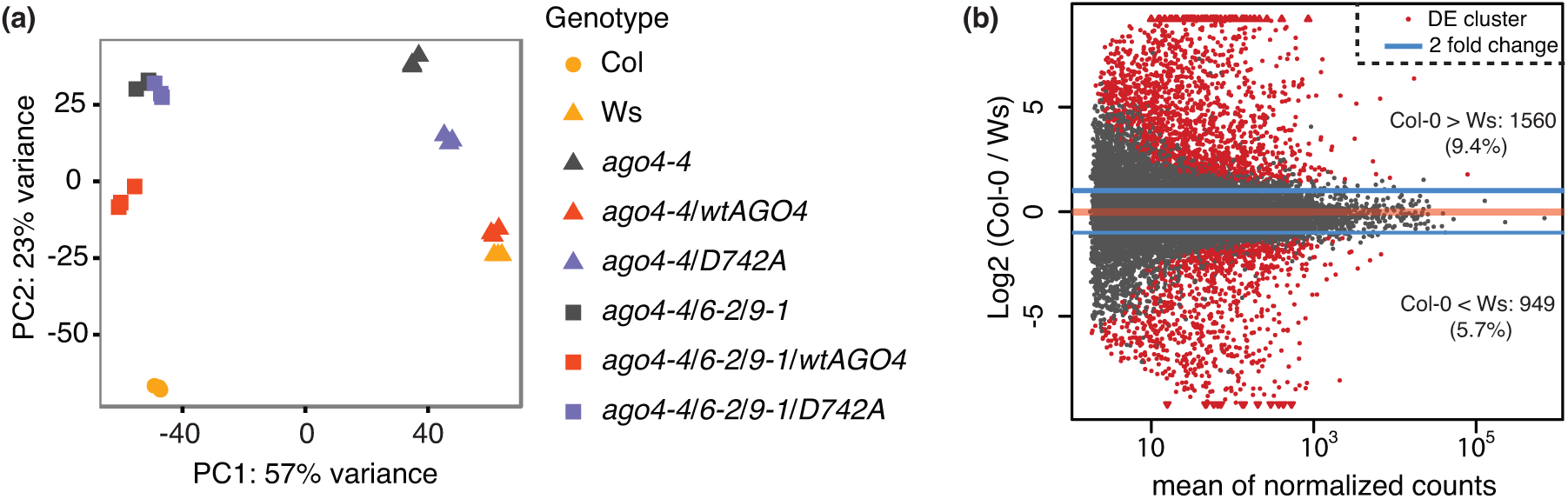
Divergence of small RNA accumulation between Col-0 and Ws. (a) Principal component analysis demonstrating overall relationships between sRNA-seq libraries. The first two principal components are shown for tested samples. (b) MA plot highlighting in red small RNA clusters with at least 2-fold differences (FDR=0.01) between Col-0 and Ws ecotypes.

## Supporting tables

**Table S1.**
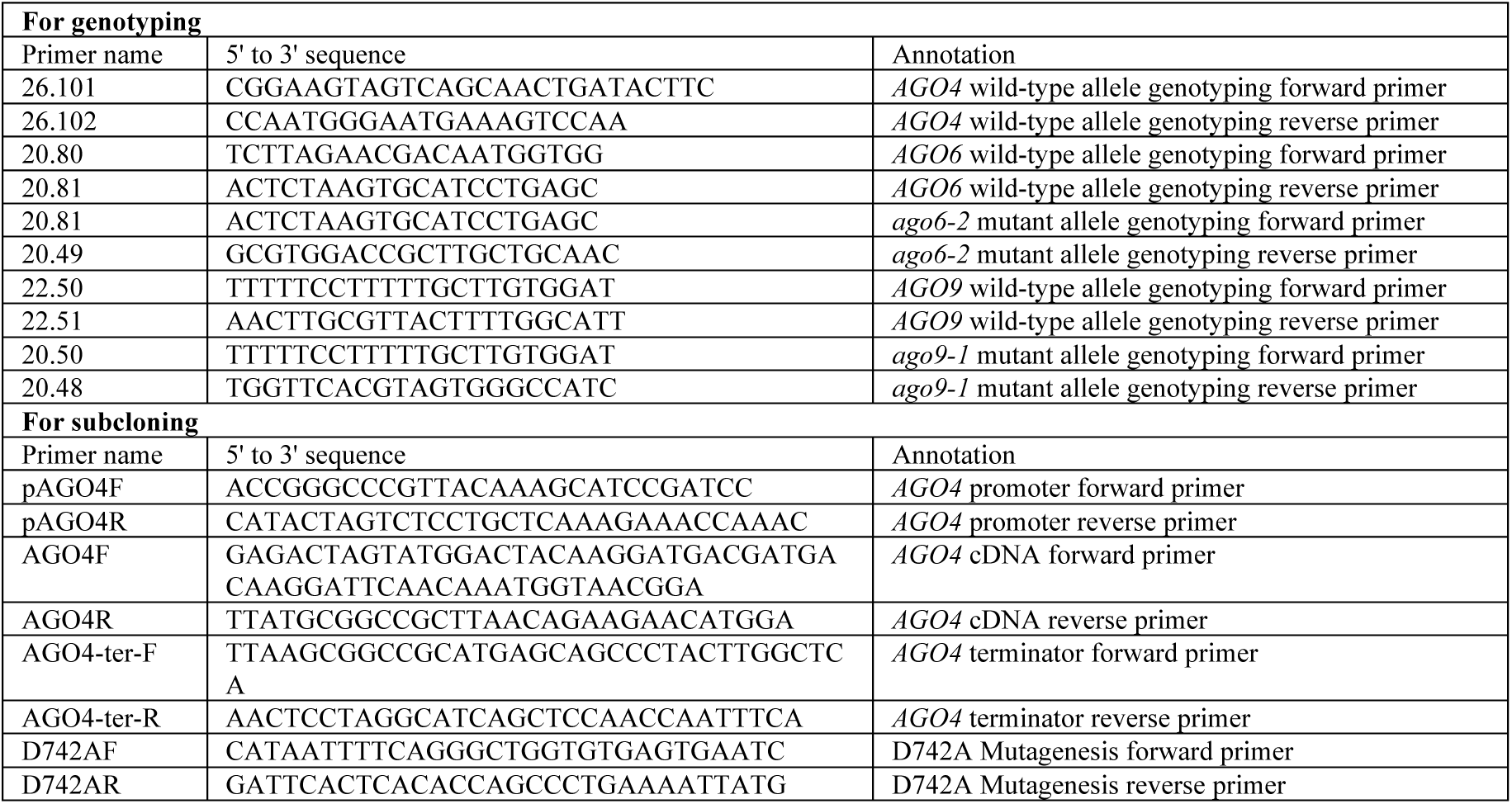
Primers used in this study.

**Table S2.** Data sources and accession numbers of *Arabidopsis thaliana* sRNA-seq libraries.

| Accession number | Genotype | Ecotype | 3'-Adapter (first 8 nt) | Source |
| --- | --- | --- | --- | --- |
| GSM2086247 | Col-0 | Col-0 | TGGAATTC | This study |
| GSM2086248 | Col-0 | Col-0 | TGGAATTC | This study |
| GSM2086249 | Col-0 | Col-0 | TGGAATTC | This study |
| GSM2327936 | Ws | Ws | TGGAATTC | This study |
| GSM2327937 | Ws | Ws | TGGAATTC | This study |
| GSM2327938 | Ws | Ws | TGGAATTC | This study |
| GSM2327939 | *ago4-4* | Ws | TGGAATTC | This study |
| GSM2327940 | *ago4-4* | Ws | TGGAATTC | This study |
| GSM2327941 | *ago4-4* | Ws | TGGAATTC | This study |
| GSM2327945 | *ago4-4/wtAGO4* | Ws | TGGAATTC | This study |
| GSM2327946 | *ago4-4/wtAGO4* | Ws | TGGAATTC | This study |
| GSM2327947 | *ago4-4/wtAGO4* | Ws | TGGAATTC | This study |
| GSM2327948 | *ago4-4/D742A* | Ws | TGGAATTC | This study |
| GSM2327949 | *ago4-4/D742A* | Ws | TGGAATTC | This study |
| GSM2327950 | *ago4-4/D742A* | Ws | TGGAATTC | This study |
| GSM2327942 | *ago4-4/6-2/9-1* | Col-0/Ws | TGGAATTC | This study |
| GSM2327943 | *ago4-4/6-2/9-1* | Col-0/Ws | TGGAATTC | This study |
| GSM2327944 | *ago4-4/6-2/9-1* | Col-0/Ws | TGGAATTC | This study |
| GSM2327951 | *ago4-4/6-2/9-1/wtAGO4* | Col-0/Ws | TGGAATTC | This study |
| GSM2327952 | *ago4-4/6-2/9-1/wtAGO4* | Col-0/Ws | TGGAATTC | This study |
| GSM2327953 | *ago4-4/6-2/9-1/wtAGO4* | Col-0/Ws | TGGAATTC | This study |
| GSM2327954 | *ago4-4/6-2/9-1/D742A* | Col-0/Ws | TGGAATTC | This study |
| GSM2327955 | *ago4-4/6-2/9-1/D742A* | Col-0/Ws | TGGAATTC | This study |
| GSM2327956 | *ago4-4/6-2/9-1/D742A* | Col-0/Ws | TGGAATTC | This study |
| GSM1103235 | Col-0 | Col-0 | TGGAATTC | Law *et al.*, 2013 |
| GSM1103237 | *nrpd1-4* | Col-0 | TGGAATTC | Law *et al.*, 2013 |
| GSM1103238 | *nrpe1-4* | Col-0 | TGGAATTC | Law *et al.*, 2013 |
| GSM1103240 | *drm2-2* | Col-0 | TGGAATTC | Law *et al.*, 2013 |
| GSM1103239 | *shh1-1* | Col-0 | TGGAATTC | Law *et al.*, 2013 |
| GSM893112 | Col-0 | Col-0 | CACTCGGG | Lee *et al.*, 2012 |
| GSM893113 | Col-0 | Col-0 | CACTCGGG | Lee *et al.*, 2012 |
| GSM893114 | Col-0 | Col-0 | CACTCGGG | Lee *et al.*, 2012 |
| GSM893115 | *nrpe1-1* | Col-0 | CACTCGGG | Lee *et al.*, 2012 |
| GSM893116 | *nrpe1-1* | Col-0 | CACTCGGG | Lee *et al.*, 2012 |
| GSM893117 | *nrpe1-1* | Col-0 | CACTCGGG | Lee *et al.*, 2012 |

